# Modeling the rarest of the rare: A comparison between joint species distribution models, ensembles of small models, and single-species models at extremely low sample sizes

**DOI:** 10.1101/2022.06.21.497071

**Authors:** Kelley D. Erickson, Adam B. Smith

## Abstract

Determining the distribution and environmental preferences of rare species threatened by global change has long been a focus of conservation. Typical minimum suggested number of occurrences ranges from ∼5 to 30, but many species are represented by even fewer occurrences. However, several newer methods may be able to accommodate such low samples sizes. These include Bayesian joint species distribution models (JSDMs) which allow rare species to statistically “borrow strength” from more common species with similar niches, and ensembles of small models (ESMs), which reduce the number of parameters by averaging smaller models. Here we explore how niche breadth and niche position relative to other species influence model performance at low sample sizes (N=2, 4, 8, 16, 32, 64) using virtual species within a community of real species. ESMs were better at discrimination tasks for most species, and yielded better-than-random accuracy even for N=2. In contrast, “traditional” single species or JSDMs were better able to estimate the underlying response curves of variables that influenced the niche, but at low sample sizes also were more likely to incorrectly identify unimportant factors as influential. Species with niches that were narrow and peripheral to the available environmental space yielded models with better discrimination capacity than species with broad niches or niches that were similar to those of other species, regardless of whether the modeling algorithm allowed for borrowing of strength. Our study suggests that some rare species may be able to be modeled reliably at very low sample sizes, although the best algorithm depends on number of occurrences and whether the niche or distribution is the focus.

---

In the face of unprecedented threats to biodiversity, understanding species’ niche preferences and distributions is as critical for conservation as ever (Guisan et al. 2013). Species distribution models (SDMs), which correlate species’ presences with environmental conditions, are widely used both to determine the current and past distribution of species as well as to predict how species will respond to global change (Elith and Leathwick 2009). However, many critically endangered species have very few occurrence records (Lomba et al. 2010, Zizka et al. 2018) and low sample sizes have been identified as one of the factors most often found to reduce model accuracy (Stockwell and Peterson 2002, Hernandez et al. 2006, Wisz et al. 2008, Thibaud et al. 2014, Santini et al. 2021, Butler and Sanderson 2022). Suggested minimum number of occurrences range from approximately 5 to greater than 200 for “traditional” single-species SDMs (Wisz et al. 2008, van Proosdij et al. 2016, Santini et al. 2021). Sample size is often only somewhat under a modeler’s control, either due to the inherent scarcity of the species or due to limited resources or difficulty in sampling. Given the abundance of species with very little data and the urgency of understanding their niche preferences and distributions, how should modelers working with the rarest of the rare proceed (Yoccoz 2022)?

Traditional modeling methods (e.g., single-species generalized linear models) typically perform poorly for species with few occurrences (Wisz et al. 2008, Santini et al. 2021). A general rule-of-thumb is to include ∼10 occurrences per coefficient, which limits the number of covariates and complexity of terms allowable in a model when sample size is low (Brun et al. 2020). However, for species with few occurrences, this can severely limit the complexity and thus the realism of models that can be constructed. Low sample size also increases model variance (uncertainty in coefficient estimates or sensitivity of coefficient estimates to small changes in the data) and bias (mis-estimation of coefficients), and risks inducing collinearity between predictors, which can compound variance and bias (James et al. 2013, Winship and Western 2016). These issues are further exacerbated when models are mis-specified (e.g., inclusion of unimportant predictors; James et al. 2013, Winship and Western 2016).

Several newer techniques hold promise for modeling species with few occurrences. Joint species distribution models (JSDMs), where species within a community are modeled simultaneously, have potential for improved modeling of rare species (Ovaskainen et al. 2010, Ovaskainen and Soininen 2011, D’Amen et al. 2017). Specifically, JSDMs assume that the coefficients that define the environmental responses of each species in the community share a common statistical distribution. This allows rare species to “borrow strength” from their more common counterparts and thus enables inclusion of species that otherwise may have too few occurrences for application of traditional, single-species model (Zipkin et al. 2009, Ovaskainen and Soininen 2011, Tingley and Beissinger 2013, Hui et al. 2013, Zhang et al. 2020). JSDMs thus assume that rare species’ environmental responses mirror those of more common species.

Another promising method employs ensembles of small model (ESMs), in which multiple simple models with just a few terms are fit and then weighted by model performance to assemble an ensemble model (Lomba et al. 2010, Breiner et al. 2015). By fitting simple models one-by-one, ESMs address the increased variance inherent in fitting multiple coefficients with sparse data, but may not be able to address biases inherent in how small samples represent environmental space. ESMs are more accurate than traditional species distribution models, especially as sample size decreases, although systematic assessments to date have only explored sample sizes ranging down to 10 (Breiner et al. 2015). A recent suggests ESMs, and JSDMs especially, are promising avenues for modeling of rare species (Jeliazkov et al. 2022).

Species’ niche breadth and position can also influence model performance, though this may interact with the modeling algorithm. Species that prefer a restricted set of environments and/or environments that are peripheral to the sampled environmental space have niches that tend to be easier for models to differentiate from available environmental space (Santika 2011, Smith et al. 2013, Soultan and Safi 2017, Connor et al. 2018). However, the borrowing of strength inherent in joint-species approaches tends to draw responses of rare species towards those of more common species (Zipkin et al. 2009, Clark et al. 2014). Hence, the ability of JSDMs to model rare species may be more affected by how similar rare species’ niches are to those of common species. In these cases, the breadth and position of rare species’ niches may be less important than their similarity to other species’ environmental preferences.

Determining the best approach to modeling very rare species depends on whether the goal is to predict the geographical distribution of the species (species distribution modeling) or to estimate the fundamental niche (ecological niche modeling; Peterson et al. 2011, Guillera-Arroita et al. 2015). For example, models can be used to locate previously unknown populations (Guisan et al. 2006, Le Lay et al. 2010) or as a means to estimate species’ current distributions for identifying areas of conservation concern (e.g., Carroll et al. 2010). In both situations, the focus is on determining the geographic distribution of the species, so discrimination (ability to differentiate between presence and absence) is important (Norberg et al. 2019). Alternatively, modelers may be more interested in estimating species’ environmental tolerances, for say, identifying suitable habitat for establishing new populations of a species (e.g., Albrecht and Long 2019), or predicting how a species may respond to a changing climate (Thuiller et al. 2006). In these cases, prioritizing model calibration (ability to identify suitable environments, regardless of whether a species occurs there) is more important (Norberg et al. 2019).

Here, we use virtual species, simulated on a real landscape within a community of real species, to determine how number of occurrences and its interaction with niche breadth and position influence model performance for ESMs, JSDMs, and “traditional” single-species distribution models (SSDMs). Our goal was to identify potential scenarios in which certain models would perform better than the rest. Specifically, we hypothesized that borrowing of strength would allow JSDMs to out-perform SSDMs and ESMs when the focal species was very rare yet similar in its niche position to the other species in the community. However, as sample size increased or the species’ niche became more distinct from that of the other species’, we expected SSDMs and especially ESMs to perform better (Smith et al. 2013, Breiner et al. 2015, Connor et al. 2018).

## MATERIAL AND METHODS

### Study System

The data component of our study is derived from Norberg et al. (2019), who found that JSDMs generally out-performed SSDMs for real communities. We used the tree community data from US Forest Service’s Forest Inventory and Analysis (as prepared by Norberg et al. 2019; https://zenodo.org/record/2637812), which is comprised of 1200 plots, with presence/absence data for 63 species. The original FIA dataset had 38 environmental covariates, which Norberg et al. reduced to 3 principal components. To more realistically approximate sampling effort for a rare species, as well as to decrease computational time, we subset our data to only include the 474 plots located in the Southeastern United States (Oklahoma, Arkansas, Tennessee, North Carolina, South Carolina, Texas, Louisiana, Mississippi, Alabama, Georgia and Florida). Norberg et al. (2019) discarded species that occurred at fewer than 10 sites, as well as species that were not represented by at least one presence in each of their cross-validation training data sets. Our subsampling resulted in 8 species with fewer than 10 occurrences (Supplementary material Appendix 1 Figure S1), but we opted to retain these in the set of “real” species since inclusion of rare species can improve results from joint-species modeling (Ovaskainen and Soininen 2011) and because our focus was on very rare species.

### Simulated Species

We considered two axes along which species could differ from each other: niche breadth (defined by the standard deviation across environmental covariates determining a species’ fundamental niche) and its relative position within the environmental space occupied by the community (the community-average niche or more extreme). We thus simulated four different species types: a species with a narrow fundamental niche breadth centered on the community average (narrow/average), a species with a broad niche breadth centered on the community average (broad/average), a species with a narrow niche breadth further away from the community average (narrow/extreme), and lastly a species with a broad niche breadth further away from the community average (broad/extreme).

In preliminary modeling of the real community, we found that many of the species only responded to 1 or 2 of the environmental principal components (PCs). We therefore defined our simulated species as responding to PC1 and PC3 while being agnostic towards PC2, but provided PC2 to the SDMs to recreate the likely common case where a modeler is only partially knowledgeable about the factors driving species’ distributions.

We simulated the species’ fundamental niches using a bivariate normal distribution, normalized so the maximum probability of presence across sites was 1 (Meynard et al. 2019), using the dmvnorm() function in the R package mvtnorm (Genz and Bretz 2009, Genz et al. 2021). We defined a broad niche breadth as having a standard deviation equivalent to the 90^th^ quantile of niche breadths across the real species in each of the two niche axes. Species with an average niche had their optimum located at the average of the mean environment occupied across species (Fig. 1). To choose the extreme and narrow niche breadth values, we explored a range of means and standard deviations. We chose a combination where the suitability function still had a relatively high probability of occurrence across the most suitable 128 sites to ensure that there was enough variation between replicates. Our narrow-niched species thus had a standard deviation that fell below the 4.8th and 69.4th quantiles of standard deviations for PCs 1 and 3, respectively, across all real species (Fig. 1).

**Figure 1.**
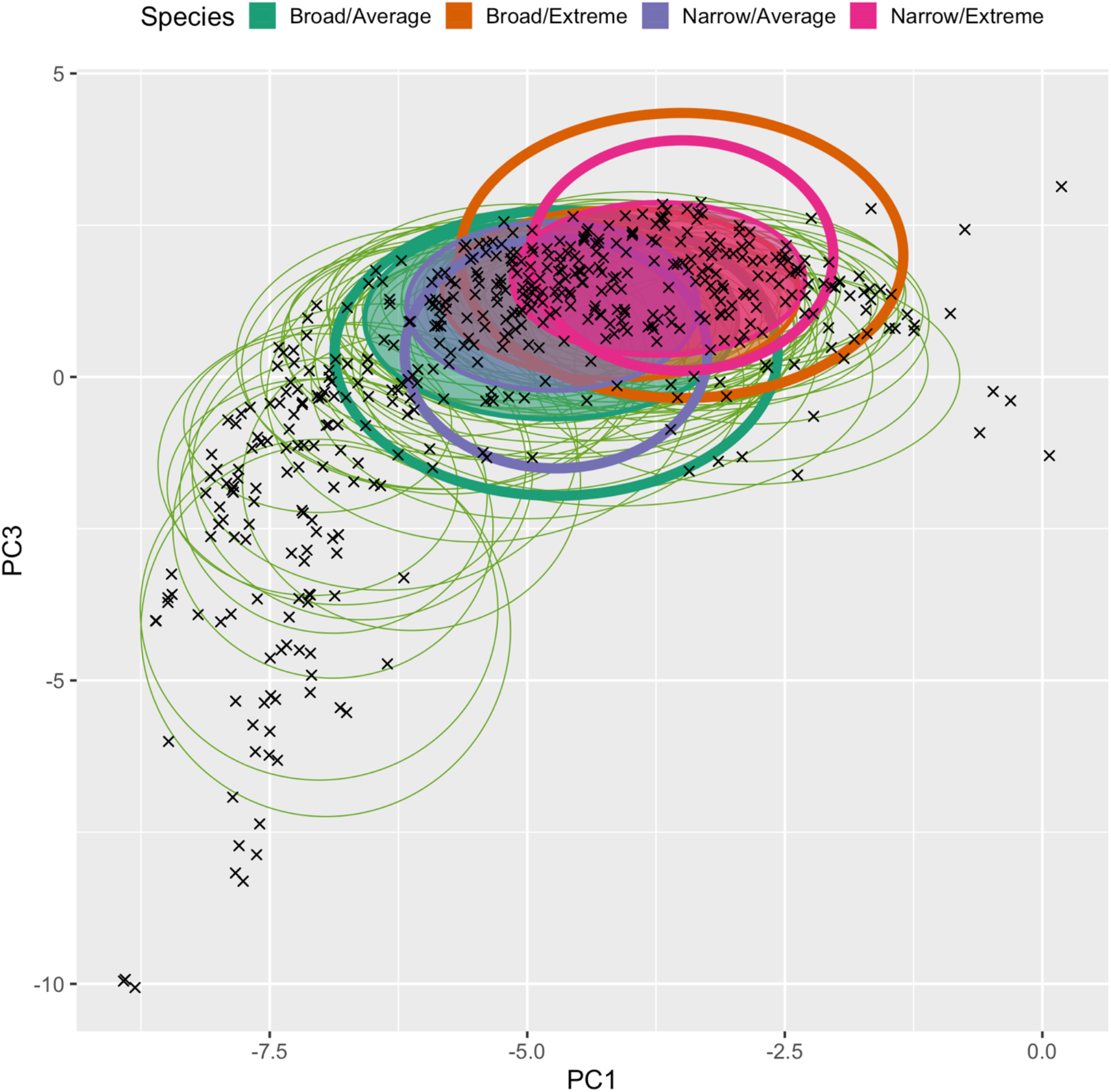
Real and simulated species in environmental space. Thin ovals represent the inner real species’ realized niches and are centered on the mean occupied environment, with axes equal to ±1 sd. Thick ovals represent fundamental and realized niches for the four types of simulated species used in the study: a broad niche centered on the mean environmental conditions at sites where species are present (broad/average), a narrow niche centered on the community average environment (narrow/average), a broad niche centered on a point in environmental space more extreme than the community average (broad/extreme), and a narrow niche centered on the extreme point (narrow/extreme). Each simulated species has two ovals, one indicating fundamental niche breadth and position (open ovals), and one the average realized niche breadth and position given the arrangement of sites (shaded ovals). Realized niches for simulated species were calculated from the mean and standard deviation of 1000 iterations of choosing N=64 presences. Colorblind-friendly colors from www.ColorBrewer.org by Cynthia A. Brewer, Geography, Pennsylvania State University.

For each combination of niche type and sample size we generated 30 replicates of *2N* presences according to the bivariate habitat suitability function, where *N* was in {2, 4, 8, 16, 32, 64}. For each replicate, we split the data into two folds such that each fold had *N* presences. SDMs were calibrated on one set and evaluated against the other, after which the sets were swapped, and the models trained on the second set and evaluated against the first. This was repeated three times for three different splits for a given set of occurrences. We ensured that there was sufficient variation across replicates among species with the same niche type and sample size (Meynard and Kaplan 2013) by constructing a histogram of the site IDs that were occupied and checked that there was variation among the 30 replicates.

### Single Species SDM

We used a single-species Bayesian generalized linear models implemented in the Hmsc package (Tikhonov et al. 2020) in R version 4.1.1 (R Core Team 2021) with 25000 iterations after a burn-in of 2000, with default priors. Models included linear and quadratic terms for each predictor. We used the default priors in the Hmsc package.

### Joint Species SDM

We implemented JSDMs again using the Hmsc package (Tikhonov et al. 2020) with 10000 iterations after a burn-in of 2000 with default priors. Models included linear and quadratic terms for each predictor. Again, we used the default priors. We focus on the HMSC implementation because Norberg et al. (2019) found it generally outperformed other joint-species and single-species modeling frameworks in predictive ability at both the community and individual species level (Norberg et al. 2019).

### Ensembles of Small Models

As presented in Lomba et al. (2010) and Breiner et al. (2015), ESMs comprise an ensemble of small models each with just two predictors, each with linear terms. Using the glm function in the stats package (R Core Team 2021), we constructed two variants of ESMs, a “simple” type using just one or two predictors regardless of sample size, and a “complex” type where we considered a total of up to 26 sub-models depending on the number of presences (model types listed in Supplementary Material Appendix S1 Table 1). The most complex models included linear and quadratic terms for each predictor. Somer’s D (2 *AUC* − 1) was used to assign weights to sub-models, with models with a D < 0 assigned a weight of zero (Breiner et al. 2015).

### Model convergence

For the SSDMs and JSDM we considered a model to have converged if the Gelman-Rubin statistic 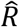 was <1.2 (Gelman and Rubin 1992). For ESMs, we considered a model to have not “converged” when all sub-model weights were <0.

### Evaluating model performance

To evaluate the ability of models to discriminate between sites where the species was present and absent, we calculated AUC; Tjur’s R^2^ (the average model prediction at presence sites minus the average prediction at absence sites; Tjur 2009); and the area under the precision-recall curve (AUC_pr_; Keilwagen et al. 2014, Grau et al. 2015, Sofaer et al. 2019). AUC indicates the probability a randomly-chosen occurrence has a higher model prediction than a random absence (Swets 1988, Mason and Graham 2002), Tjur’s R^2^ indicates the ease of discriminating between presences and absences sites given model predictions, and AUC_pr_ is less sensitive to class imbalance than AUC (Sofaer et al. 2019).

To evaluate calibration accuracy, we measured the match between the species’ actual response curves from the generative niche model and estimated responses from the SDMs. For PC1 and PC3, the environmental covariates which were used to generate the simulated species, we used the compareResponse function in the R package enmSdm (Smith 2021) to calculate the rank correlation between the actual habitat suitability response curves and the models’ predicted response curve across the environmental range of all sites. Models that are better able to recover the species response curve should have higher rank correlation. For PC2, which was not used to generate the simulated species, we calculated the standard deviation of the models’ predicted response curves as PC2 was varied from the minimum to the maximum value across all plots while values of PCs 1 and 3 were held constant at the mean value across the species’ occurrences. Well-calibrated models should have a standard deviation of zero, as they should predict a flat response of the species probability of presence to PC2. We also evaluated overall model accuracy using a weighted root-mean square error (RMSE):

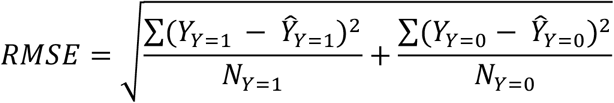

where Y is the true presence (Y=1) or absence (Y=0) of a site, *Ŷ* is the predicted probability of presence and N is the number of sites where the species is present (Y=1) or absent (Y=0).

For each combination of niche type and sample-size replicate, we averaged each performance metric over the two folds within each split and then across folds. We did not conduct statistical hypothesis tests because we know, a priori, that differences exist between species and sample sizes (White et al. 2014). Rather, we focused on effect size differences by examining overlap between the inner 80^th^ percentile of values of each test statistic (AUC, R^2^, AUC_pr_, RMSE, and rank correlation) across species and sample size (White et al. 2014).

All analyses were done in R version 4.1.1 (R Core Team 2021) Analyses relied primarily on the packages Hmsc(Tikhonov et al. 2020), enmSdm (Smith 2021), and PRROC (Keilwagen et al. 2014, Grau et al. 2015) packages. Code for running all analyses is available at https://github.com/kerickson22/SDMs_for_rare_species_modeling and data for the real species from https://zenodo.org/record/2637812 (Norberg et al. 2019).

## RESULTS

### Model Convergence

ESMs converged more frequently than either the SSDMs and JSDMs (Figure 2). Both SSDMs and JSDMs converged in roughly half of all cases regardless of sample size. Running more scenarios would have provided more models that converged for analysis, but in real-world situations non-convergence is affected by properties of the data (in addition to other factors; Gelman et al. 2013). Since modelers are often not in a position to acquire more data easily, and because we ran our models in the same manner as used in other studies (Breiner et al. 2015; Norberg et al. 2019), we decided to retain only the models that did converge, although there was little qualitative difference between using all models instead of just those that converged (Supplementary Material Appendix 1 Figures S2-S5).

**Figure 2:**
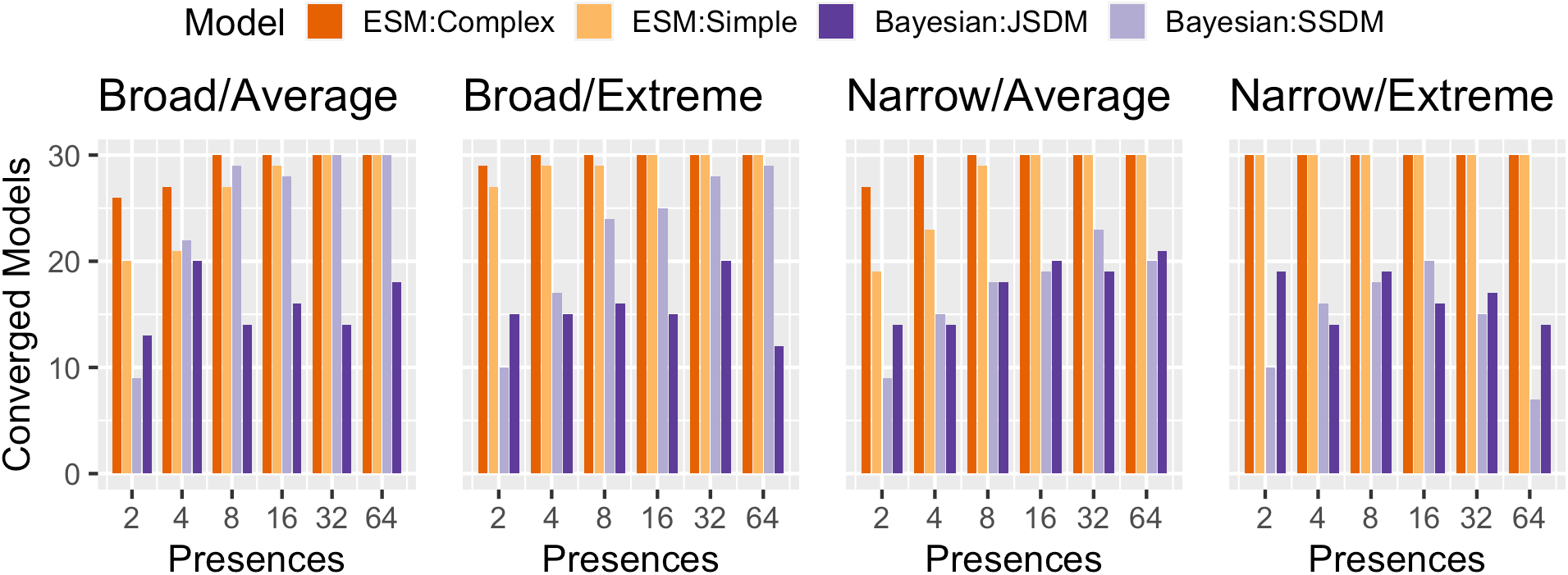
Number of converged models as a function of species’ niche breadth and position, and number of calibration occurrences. Ensembles of small models (complex and simple) were considered to be converged if at least one sub-model had a nonzero weight. Bayesian models (joint and single-species) were considered converged if all values of their Gelman-Rubin statistic 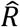 were < 1.2. Colorblind-friendly colors from www.ColorBrewer.org by Cynthia A. Brewer, Geography, Pennsylvania State University.

### Model Discrimination

Simple and complex ESMs had higher median AUC values than either of the Bayesian models, although as the number of presences increased the Bayesian models improved, though they were never better the ESMs (Figure 3a). The effect of the number of presences on simple and complex ESMs depended on the species’ niche. For the broad/average species, increasing the number of presences actually decreased median AUC for ESMs (Fig. 3a), while for the broad/extreme and narrow/extreme species it remained roughly constant of increased slightly. For the narrow/average species, increasing the number of presences increased AUC of the complex ESM while AUC of simple ESMs remained decreased slightly. SSDMs and JSDMs performed very similarly to one another. For SSDMs and JSDMs, median AUC at N = 2 was ∼0.5 for except the narrow/extreme species, which was ∼0.8 (but with inner 80^th^-quantiles encompassing or very close to 0). Increasing the number of presences improved AUC of both Bayesian models similarly. For a given number of presences, Bayesian models of species with narrow niches had higher median AUC than corresponding species with broad niches. For the same number of presences and all model types, species with extreme niches had slightly better performance than species located closer to the mean community centroid (Figure 3a).

**Figure 3.**
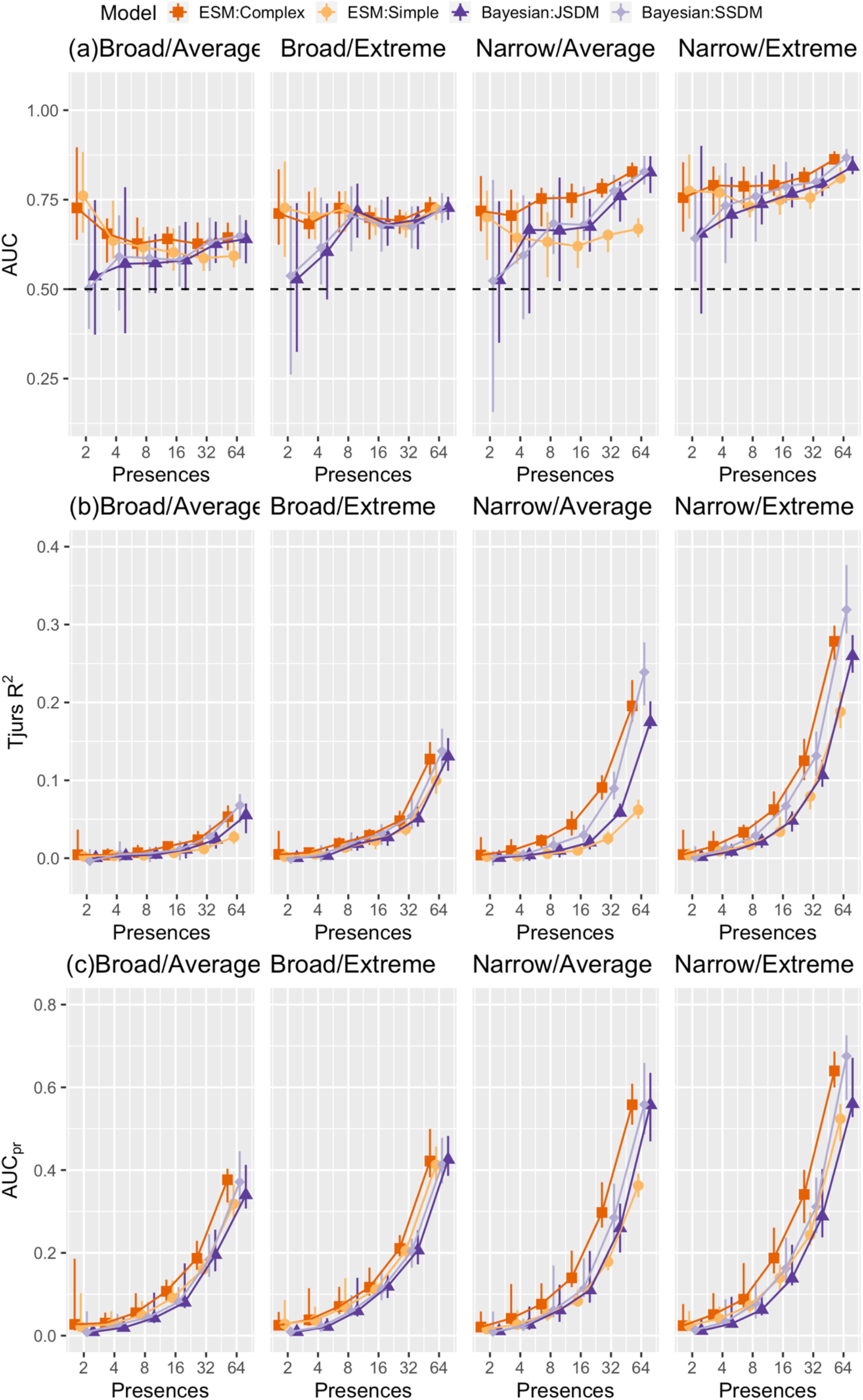
Measures of model discrimination (a) AUC, (b) Tjur’s R^2^, and (c) AUC_pr_ as a function of number of presences and species’ niche breadth and position for single-species and joint Bayesian species distribution models, and simple and complex ensembles of small models. Trend lines are included to make it easier to discern the relative position of model-types, but because the number of presences may affect the maximum (AUC) or minimum (AUC_pr_) possible value of these performance metrics comparison across different numbers of presences should be done with this in mind. Error bars represent the 90^th^ and 10^th^ quantiles across converged models. Colorblind-friendly colors from www.ColorBrewer.org by Cynthia A. Brewer, Geography, Pennsylvania State University.

For small numbers of presences, Tjur’s R^2^ and AUC_pr_ were very similar, and near 0, for all models and species (Fig. 3b and c). At larger numbers of presences (N ≥∼8-16), species with narrow niches tended to have higher R^2^ and AUC_pr_ than those with broad niches (Figure 3b). For broad-niched species (all numbers of presences) and narrow-niched species at N ≤16, there was little difference between algorithms. For narrow-niched species at larger number of presences (N > 32), SSDMs tended to have the highest R^2^ and AUC_pr_, and simple ESMs the lowest.

### Model Calibration

#### Response Curves

For all models, species, and predictors, the rank correlation between actual and estimated suitability for the two “true” variables (PCs 1 and 3) was on average fairly high (median *ρ* usually > 0.85; Figure 4a and c). However, some models were miscalibrated with PCs 1 and/or 3, with *ρ* < 0.5 or even <0. We consider values <0.5 “risky” because they indicate the models were unable to capture the true magnitude and sometimes direction of species’ relationships with environmental gradients. Model calibration varied with niche type as well as predictor. Generally, models that were well-calibrated with one or two variables were poorly calibrated with another, and no one model was risk-free. Simple ESMs had very poor calibration for PC1 for all species types, even at large numbers of presences. Complex ESMs had risky cases for low numbers of presences for all variables and species but improved as the number of presences increased. Both ESMs tended to be less sensitive to PC2 (the “false” predictor) than either of the Bayesian models for N≥8 (Figure 4b). JSDMs had poor calibration for PC 1 at small numbers of presences (except for the narrow/average species), and usually the poorest or second-poorest calibration with PC 2 for all numbers of presences. SSDM calibration had risky cases at low numbers of presences for PCs 1 and 2. Across species, narrow/average species tended to have the best calibration, for which only simple ESMs (all numbers of presences) and SSDMs (low number of presences) struggled.

**Figure 4:**
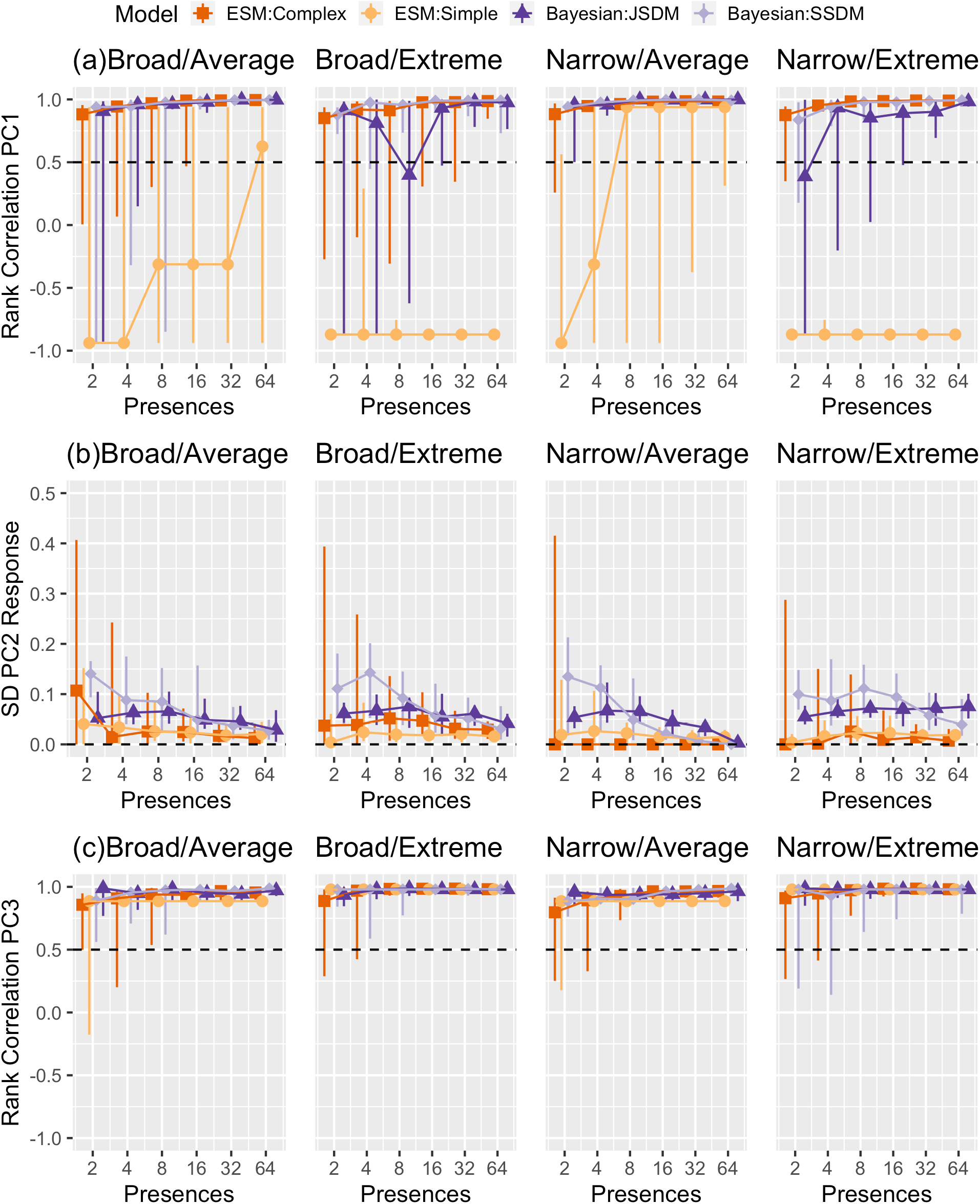
Ability of models to correctly identify response curves by species type, number of presences, and type of model. (a) Rank correlation of the real and predicted response curve for PC1, which was used to create niches. We considered values below the dashed black to indicate “risky” models incapable of accurately producing response curves. (b) Standard deviation of the predicted response curve for PC2, which was provided to models but not used to create niches. Values closer to zero indicate better calibration for PC2. (c) Rank correlation of the real and predicted response curves for PC3, which was used to create niches. Error bars encompass the 90^th^ and 10^th^ quantiles across converged models. Colorblind-friendly colors from www.ColorBrewer.org by Cynthia A. Brewer, Geography, Pennsylvania State University.

#### Weighted RMSE

For all models and all species, RMSE improved (became smaller) as the number of presences decreased (Supplementary material Appendix 1 Figure S2), but otherwise displayed trends within and across species qualitatively similar to the discrimination metrics.

## DISCUSSION

We evaluated the ability of two promising modeling methods—joint species distribution models (JSDMs) and ensembles of small models (ESMs)—for modeling species with extremely low numbers of presences. We hypothesized that at very low number of presences, species with niches that closely resembled those of other species would benefit from Bayesian joint-species modeling methods that employ “borrowing of strength”. We also hypothesized that as the number of presences increased species with narrow niches that were peripheral to the available environmental space would be modeled more accurately by ESMs because the species would have enough data to inform simple models, yet not enough to correctly parameterize a “traditional” single-species model (Lomba et al. 2010, Santika 2011, Smith et al. 2013, Breiner et al. 2015 p. 2, Soultan and Safi 2017, Connor et al. 2018). Our results depended not only on sample size, but niche characteristics and the aim of the modeling (i.e., recreating distributions versus niches) and are summarized in Fig. 5. We found that compared to methods that modeled species alone, borrowing of strength in JSDMs improved model calibration compared to a SSDM approach (Fig. 4) but not discrimination (Fig. 3). Rather, ensembles of small models (ESMs) had higher discrimination accuracy at small numbers of presences, and were less likely to misidentify unimportant variables, although they were less well-calibrated than Bayesian approaches for other parameters (Fig. 4). We also found that discrimination accuracy depended on species’ niche breadth and position in environmental space. Species with narrow niches and niches more peripheral to other species’ niches yielded the models with the highest discriminatory capacity, regardless of sample size. Niche position and niche breadth also influenced model calibration, with models for narrow-niched species centered on the community average tending to make fewer calibration mistakes than the rest of the species (Fig. 4).

**Figure 5.**
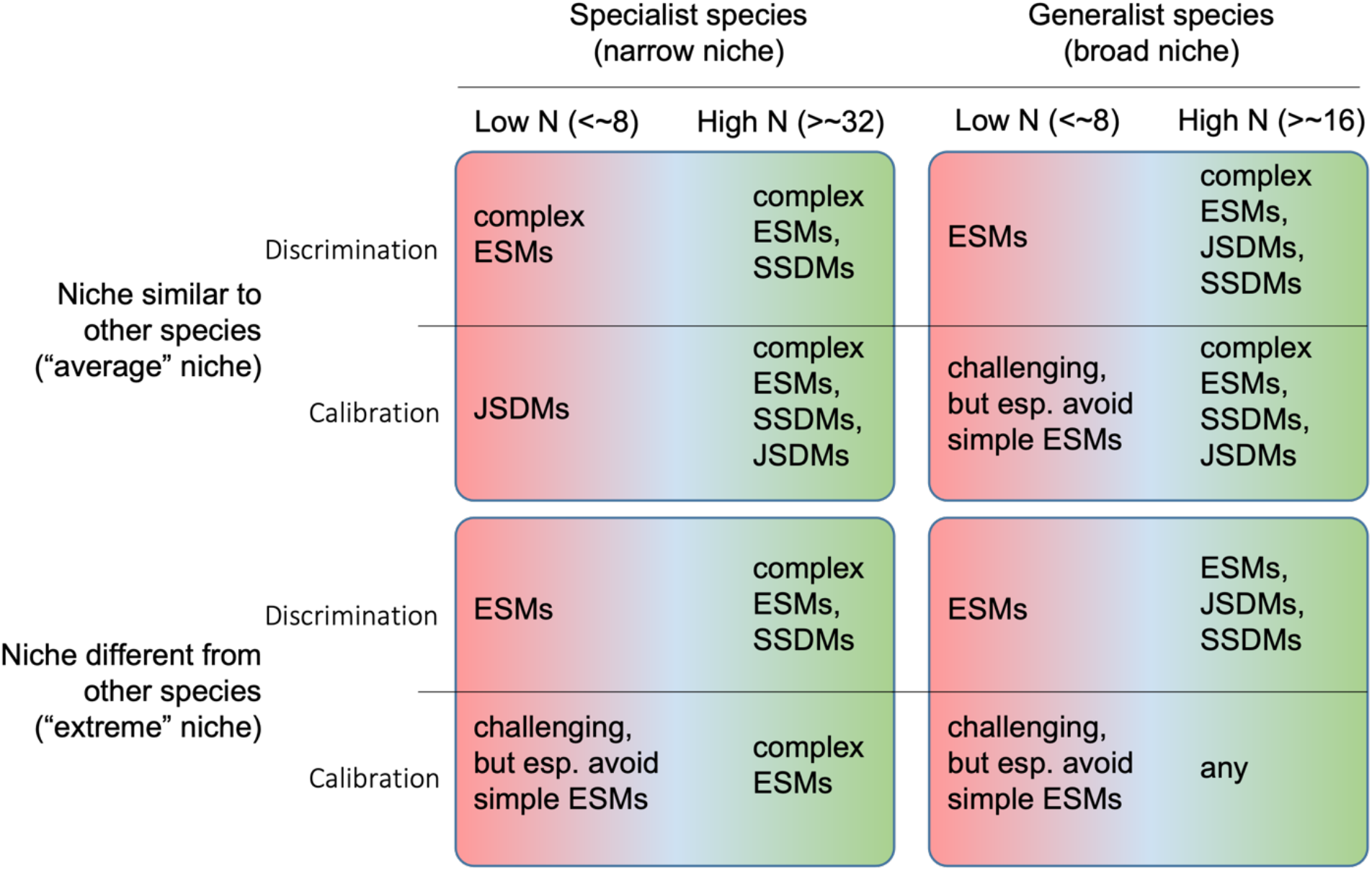
Summary of results. For each combination of modeling aim (discrimination versus calibration), niche breadth (narrow versus broad), niche position (similar to other species versus peripheral), and number of occurrences (N), we note the models that performed the best.

Determining which model type to use depends on whether the focus of the modeling effort is in understanding the fundamental niche or the geographic distribution of the species (Fig. 5). Ideally, models would be able to reproduce both a species’ geographic distribution and environmental responses reliably, but often there is a trade-off between model discrimination and calibration (Jiménez-Valverde et al. 2013). If the focus is on identifying the geographic range, ESMs would be a good choice as they had better discrimination metrics than the other models, even with the number of presences as few as N=2. However, no model type was without risk at small numbers of presences when the intended use relies on reproducing the fundamental niche (i.e., predicting how species will respond to novel environmental conditions). Simple ESMs were especially mis-calibrated, even at larger number of presences (Fig. 4). Nonetheless, increasing the number of presences generally did reduce the frequency of mis-calibrated models.

All models are subject to the bias-variance trade-off (Hastie et al. 2009). Complicated models with many parameters run the risk of overfitting the data resulting in models with low variance but high bias, while models with too few parameters underfit the data and have high variance but low bias (Hastie et al. 2009, James et al. 2013). Using frequentist-based modeling, only two parameters (without variance) can be estimated from only two presences. In the Bayesian framework however, it is possible for the number of parameters being estimated to exceed the number of datapoints while still successfully estimating these parameters, provided a suitable prior is used (Gelfand and Sahu 1999). While some of our single-species models were able to achieve convergence despite the identifiability issue at low numbers of presences with our use of vague priors (Figure 4), using more informative priors for rare species with limited data may improve model inference (Gelfand and Sahu 1999). Informative priors based on expert opinion or prior knowledge resulting from previous independent studies have been proposed as a way to improve confidence in Bayesian inference for conservation applications (Martin et al. 2005, McCarthy and Masters 2005). Data from physiological experiments or from closely related species may also be useful for inferring environmental tolerances (Qiao et al. 2017, Smith et al. 2019). As long as there is transparency in the choice of prior, using additional information in the form of an informed prior can be a way forward for making inference for particularly data-deficient species.

We found that as the number of presences decreased, ESMs, which are built from a suite of frequentist generalized linear models, performed better at discrimination (AUC) than either of the two Bayesian models. Allowing the complexity of sub-models included to increase as the number of presences yielded better results in comparison to bivariate-only (“simple”) ESMs, which sometimes even worsened with increased sample size (Fig. 3a). ESMs were least likely to falsely estimate a response to PC2 above N=8, although Bayesian models were somewhat better able to estimate the response to PC1. Simple ESMs were especially miscalibrated against PC1, despite (along with complex ESMs) having moderately reliable discrimination capacity with very few presences.

We expect that most “rare” species of conservation interest are either specialists of uncommon environments (narrow-niched; e.g. Albrecht and Long 2019) or were once common but have suffered from reduction due to anthropogenic factors (broad-niched, Channell and Lomolino 2000). Our results suggest hope especially for the former, as narrow-niched species, especially those near the margin of available environmental space, yielded the most accurate models (Fig. 4). Generalists that have suffered recent decline will, however, be especially challenging to model, especially as their current distributions likely underrepresent their environmental tolerances (Faurby and Araújo 2018). We did find that species with niches on the margin of available environmental space had models with better discrimination and calibration. Species may also be rare because environmental conditions conducive to their persistence may be rare (and thus, likely peripheral to available environmental space; e.g., Gregory and Gaston 2000, Díaz et al. 2020 but see Morueta-Holme et al. 2013). In these cases, the position of their inhabited environments may allow for more accurate modeling. However, these species may also be prone to having their apparent niche truncated because they exist on the “edge” of available environments (cf. Fig. 1). In these cases, extrapolation of models may be risky if environments to which models are projected fall “beyond” the region of truncation (Feeley and Silman 2010, Anderson 2013), so attention to model calibration will be especially important.

Our work points the way toward further potential improvements for modeling rare species. Hierarchical, joint-species Bayesian models like HMSC (Tikhonov et al. 2020) allow for inclusion of information about the underlying spatial structure of the sampling locations (e.g., Ver Hoef et al. 2018) and/or additional information about phylogenetic relationships and traits can be incorporated to improve estimates (Ovaskainen et al. 2017, Morales-Castilla et al. 2017, Tikhonov et al. 2020). For the sake of computational time available to run our full-factorial combination of models, we included only environmental predictors in the joint- and single-species hierarchical models. Test runs using JSDMs with latent factors for spatial autocorrelation required ∼8 days of run time per model for a single data instance, which we deemed unlikely to be acceptable for practical use (cf. Velásquez-Tibatá et al. 2016). However, if there is additional information about the focal species, model performance (and run time) could be improved by using a smaller subset of the total community, such as those that share functional traits or habitats with the species of interest (Pacifici et al. 2014). We used a presence-absence dataset, assuming perfect detection with no sampling bias. However, sampling bias is a common feature of occurrence data (Meyer et al. 2016). Hence, our results should be viewed as “ideal” conditions for each level of number of presences. The number of presences necessary for producing robust models in real-world situations will likely be higher, and modelers should consider additional

## CONCLUSIONS

Modelers working with rare species often must make decisions based on the limited data available. In our analysis, we show an example of a workflow (Box 1) to help determine which modeling approach to use. Plotting presences in environmental space can help determine whether a species will be easier to model (narrow niche) or more challenging (wider niche). Additionally, comparing the location of the presences in niche-space to the sampled niche-space will help identify whether the niche is peripheral to available environmental space or more interior. We found that complex ensembles of small models worked well for estimating species distributions, even at extremely low number of presences. If researchers are interested in modeling the fundamental niche however, larger numbers of presences may be necessary and Bayesian models may perform better.

**Box 1.**
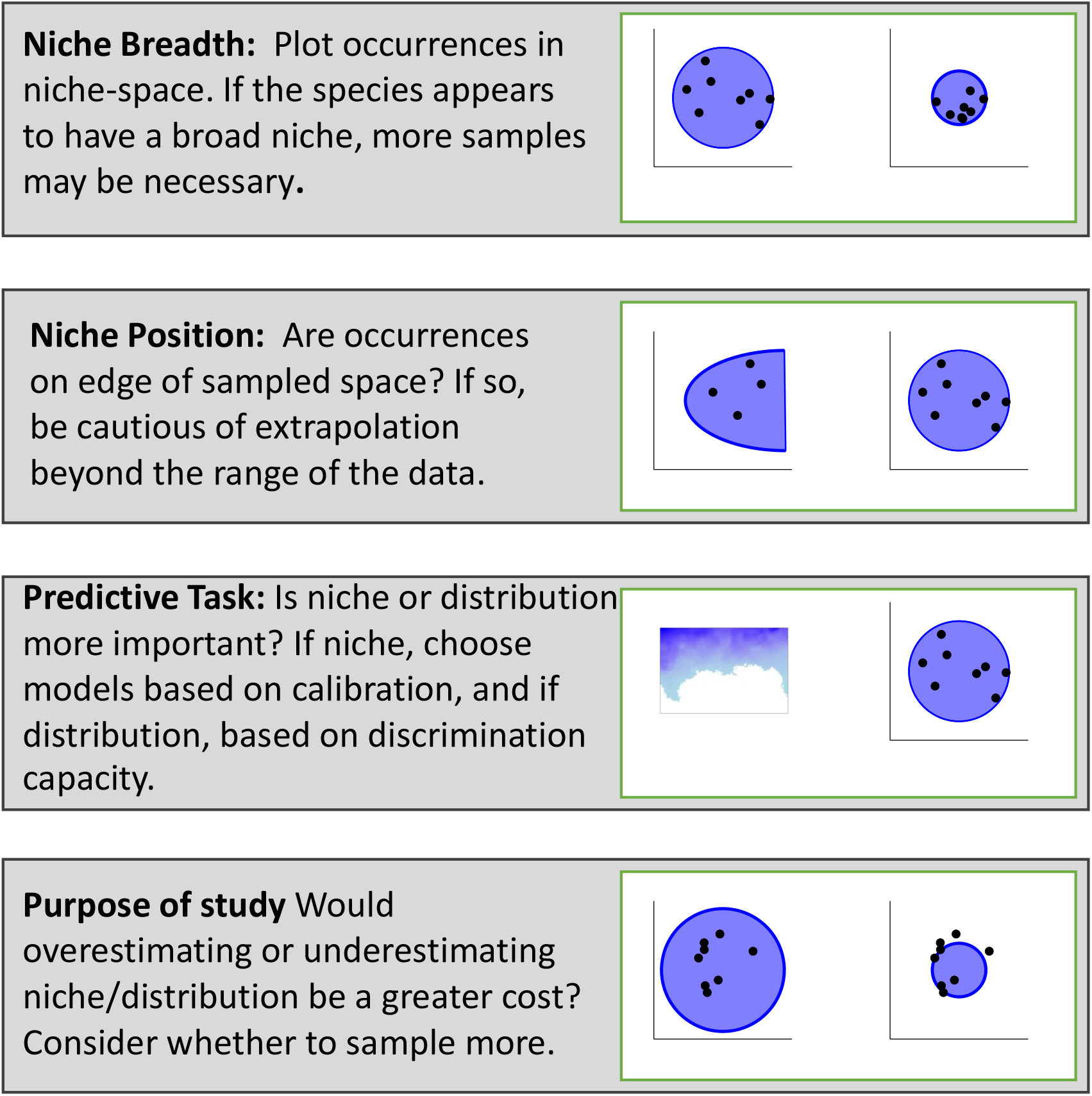
Considerations for modeling rare species.

## ACKNOWLEDGEMENTS

We thank O. Ovaskainen for answering questions about the HMSC package, and the Global Change and Conservation Lab Group at the Missouri Botanical Garden for helpful feedback on versions of this manuscript.

## DATA AVAILABILITY STATEMENT

Code for running all analyses is available at https://github.com/kerickson22/SDMs_for_rare_species_modeling, and data for the real species from https://zenodo.org/record/2637812 (Norberg et al. 2019).

