## Supplement for "Modeling the rarest of the rare: A comparison between joint species distribution models, ensembles of small models, and single-species models at extremely low sample sizes"

Table S1. Formulae included in ensembles of small models by the number of presences (N).

| | FORMULA | ESM:SIMPLE | ESM:COMPLEX |
|---|---|---|---|
| **1** | Y ~ 1 + PC1 | N ≥ 2 | N ≥ 2 |
| **2** | Y ~ 1 + PC2 | | |
| **3** | Y ~ 1 + PC3 | | |
| **4** | Y ~ 1 + PC1 + PC2 | N ≥ 4 | N ≥ 4 |
| **5** | Y ~ 1 + PC1 + PC3 | | |
| **6** | Y ~ 1 + PC2 + PC3 | | |
| **7** | Y ~ 1 + I(PC1^2) + PC1 | | |
| **8** | Y ~ 1 + I(PC2^2) + PC2 | | |
| **9** | Y ~ 1 + I(PC3^2) + PC3 | | |
| **10** | Y ~ 1 + PC1 + PC2 + PC3 | | |
| **11** | Y ~ 1 + I(PC1^2) + PC1 + PC2 | | |
| **12** | Y ~ 1 + I(PC1^2) + PC1 + PC3 | | |
| **13** | Y ~ 1 + I(PC2^2) + PC1 + PC2 | | |
| **14** | Y ~ 1 + I(PC2^2) + PC2 + PC3 | | |
| **15** | Y ~ 1 + I(PC3^2) + PC1 + PC3 | | |
| **16** | Y ~ 1 + I(PC3^2) + PC2 + PC3 | | |
| **17** | Y ~ 1 + I(PC1^2) + I(PC2^2) + PC1 + PC2 | | N ≥ 8 |
| **18** | Y ~ 1 + I(PC1^2) + I(PC3^2) + PC1 + PC3 | | |
| **19** | Y ~ 1 + I(PC2^2) + I(PC3^2) + PC2 + PC3 | | |
| **20** | Y ~ 1 + I(PC1^2) + PC1 + PC2 + PC3 | | |
| **21** | Y ~ 1 + I(PC2^2) + PC1 + PC2 + PC3 | | |
| **22** | Y ~ 1 + I(PC3^2) + PC1 + PC2 + PC3 | | |
| **23** | Y ~ 1 + I(PC1^2) + I(PC2^2) + I(PC3^2) + PC1 + PC2 + PC3 | | |
| **24** | Y ~ 1 + I(PC1^2) + I(PC2^2) + PC1 + PC2 + PC3 | | |
| **25** | Y ~ 1 + I(PC1^2) + I(PC3^2) + PC1 + PC2 + PC3 | | |
| **26** | Y ~ 1 + I(PC2^2) + I(PC3^2) + PC1 + PC2 + PC3 | | |

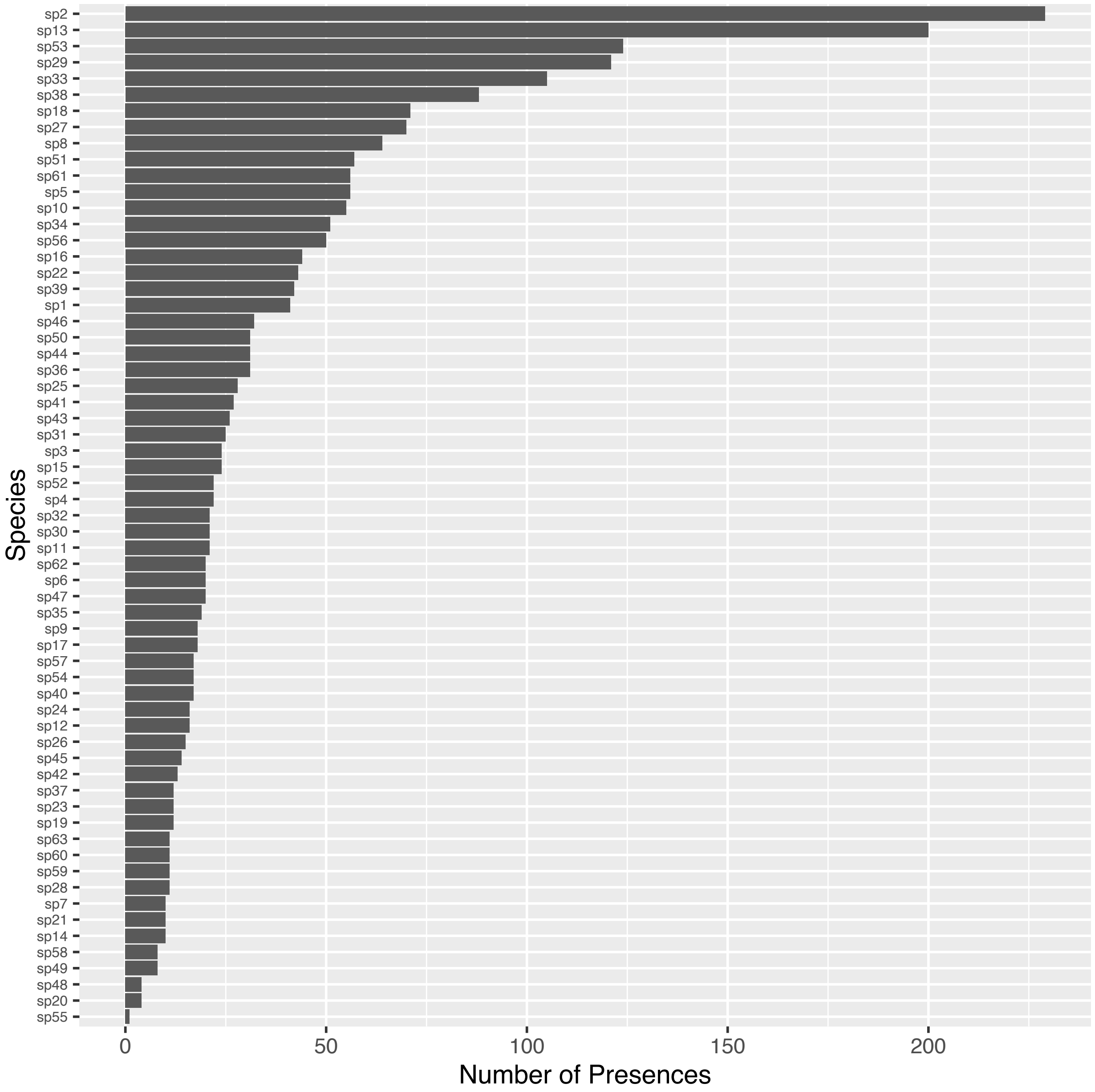


Figure S1. Number of presences within the study area for the 63 real species from Norberg et al. (2019).

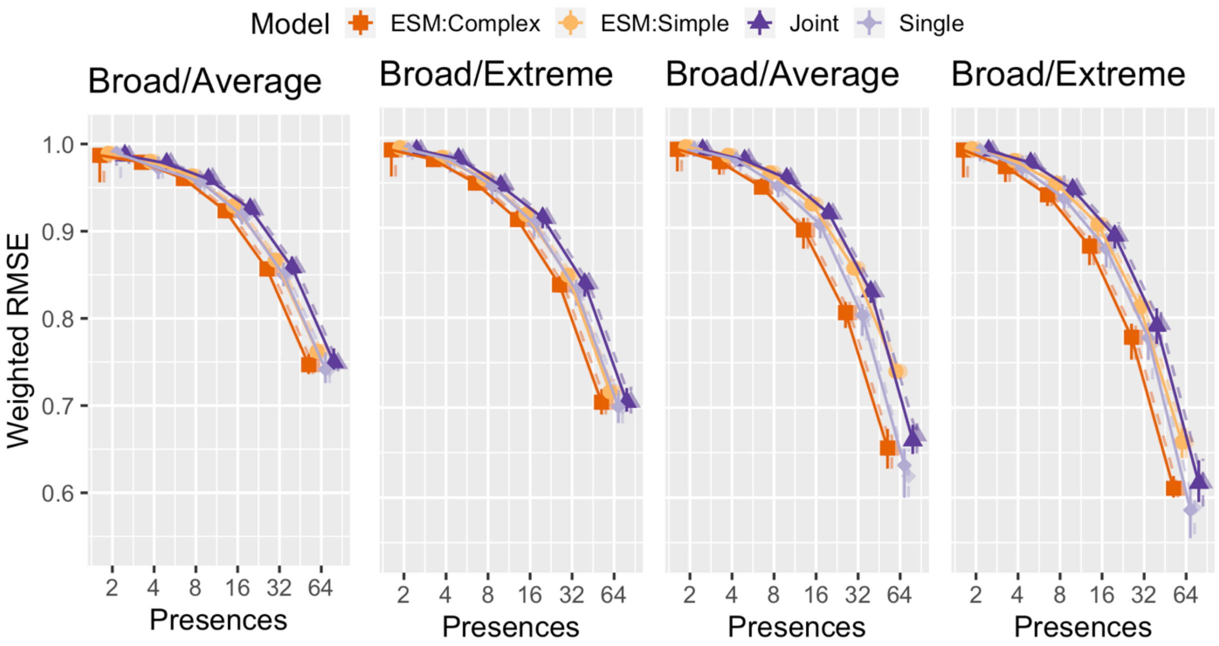


Figure S2. Weighted root-mean square error as a function of number of presences for single-species and joint Bayesian species distribution models, and simple and complex ensembles of small models. Lower values connote more accurate models. Error bars represent the 90th and 10th quantiles of values across all converged models for a given number of presences. Solid lines include only converged models, while dashed lines include all models, even those that did not converge. Colorblind-friendly colors from www.ColorBrewer.org by Cynthia A. Brewer, Geography, Pennsylvania State University.

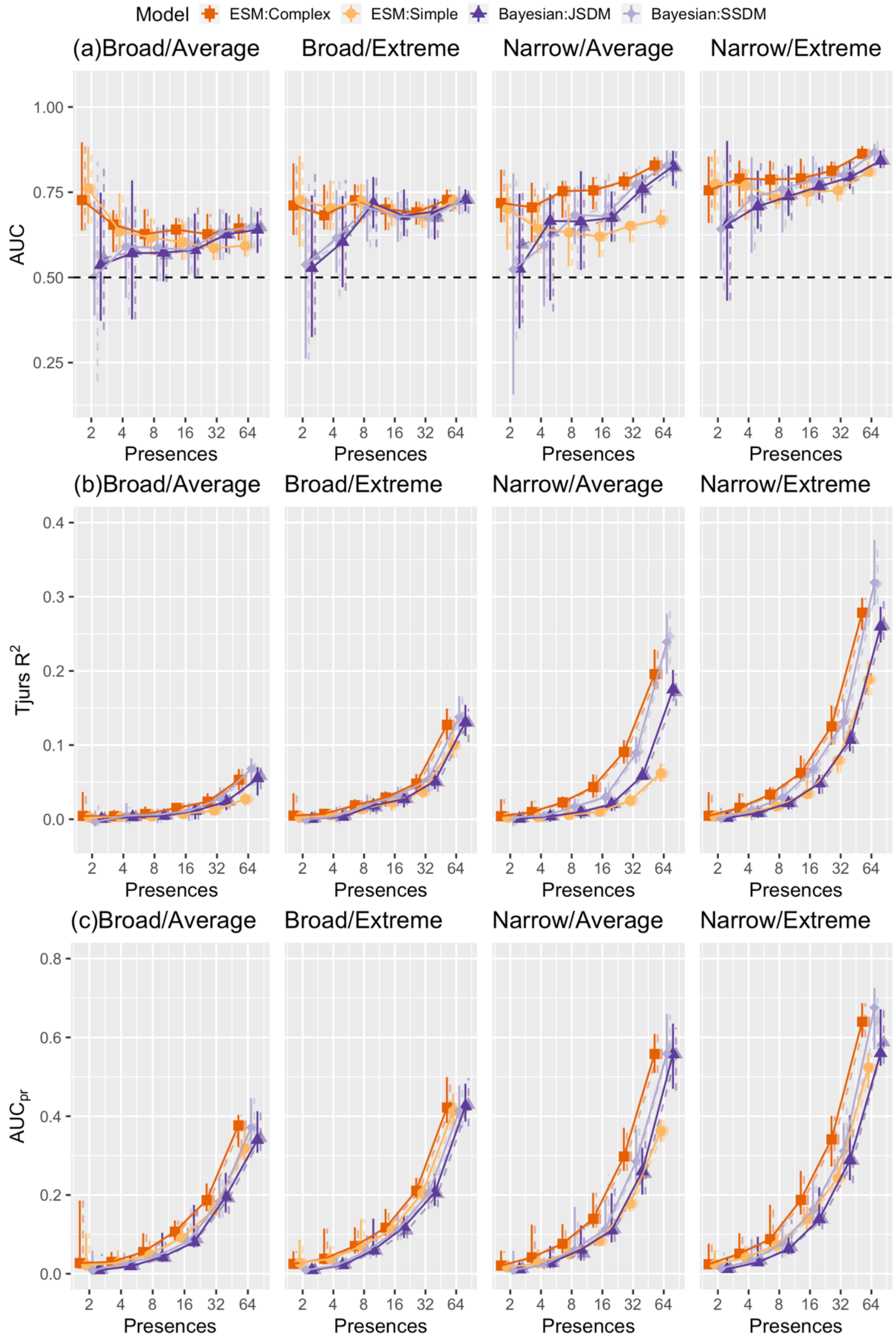
Model ESM:Complex ESM:Simple Bayesian:JSDM Bayesian:SSDM
(a)Broad/Average
Broad/Extreme
Narrow/Average
Narrow/Extreme
AUC
1.00
0.75
0.50
0.25
2 4 8 16 32 64
Presences
(b)Broad/Average
Broad/Extreme
Narrow/Average
Narrow/Extreme
Tjurs R2
0.4
0.3
0.2
0.1
0.0
(c)Broad/Average
Broad/Extreme
Narrow/Average
Narrow/Extreme
AUCpr
0.8
0.6
0.4
0.2
0.0

Figure S4. Measures of model discrimination (a) AUC, (b) Tjur's $R^2$, and (c) $AUC_{pr}$ as a function of number of presences and niche breadth and position for single-species and joint Bayesian species distribution models, and simple and complex ensembles of small models. Trend lines are included to make it easier to discern the relative position of model-types, but because the number of presences may affect the maximum (AUC) or minimum ($AUC_{pr}$) possible value of these performance metrics it is not appropriate to directly compare across different numbers of presences. Error bars represent the $90^{th}$ and $10^{th}$ quantiles. Solid lines include only converged models, while dashed lines include all models, even those that did not converge. Colorblind-friendly colors from www.ColorBrewer.org by Cynthia A. Brewer, Geography, Pennsylvania State University.

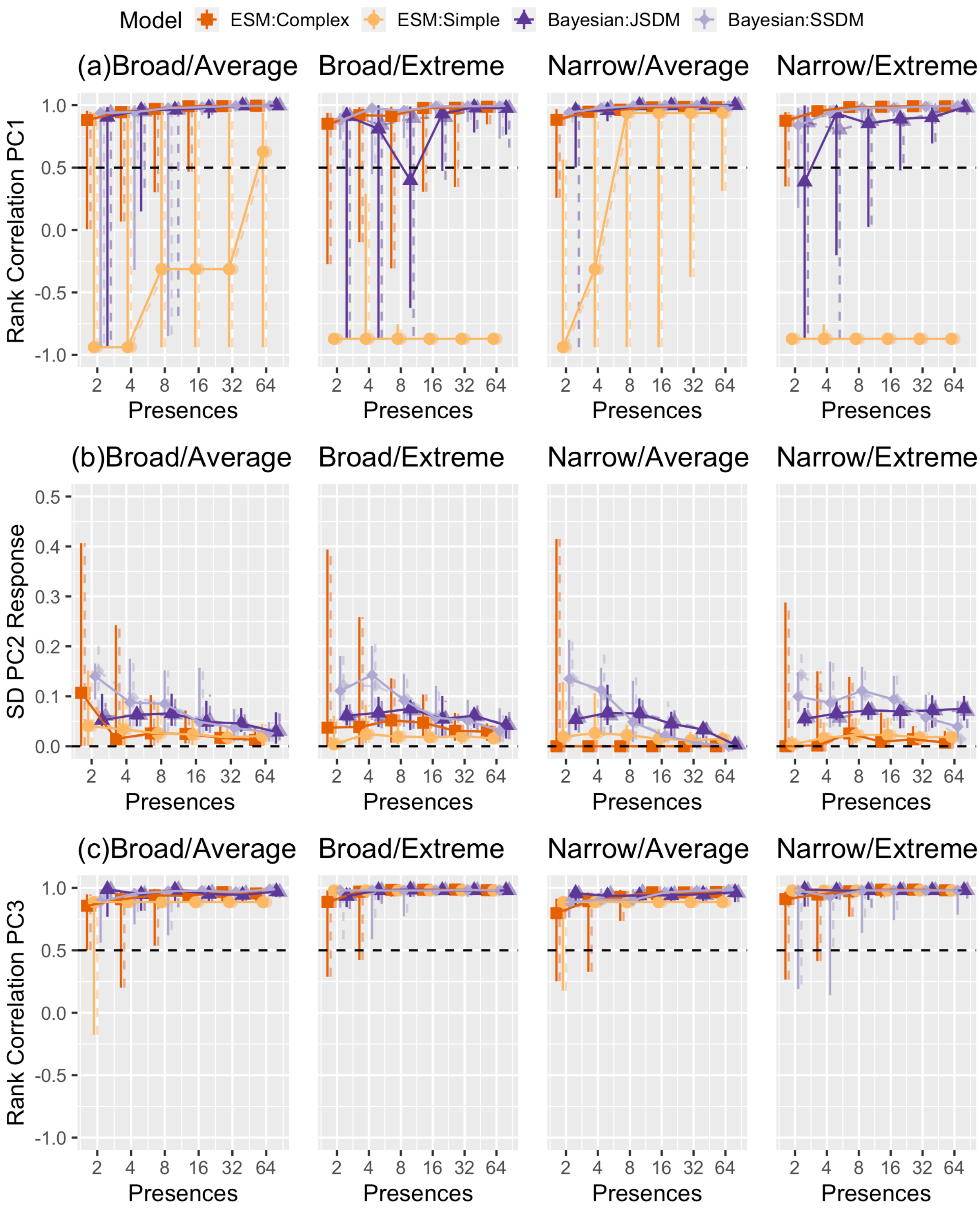


**Figure S5.** Ability of models to correctly identify response curves by species type, number of presences, and type of model. (a) Rank correlation of the real and predicted response curve for PC1, which was used to create niches. We considered values below the dashed black to indicate "risky" models incapable of accurately producing response curves. (b) Standard deviation of the predicted response curve for PC2, which was provided to models but not used to create niches. Values closer to zero indicate better calibration for PC2. (c) Rank correlation of the real and predicted response curves for PC3, which was used to create niches. Error bars encompass the 90th and 10th quantiles across converged models. . Solid lines include only converged models, while dashed lines include all models, even those that did not converge. Colorblind-friendly colors from www.ColorBrewer.org by Cynthia A. Brewer, Geography, Pennsylvania State University.
